# Frank-Starling Mechanism and Short-Term Adjustment of Cardiac Flow

**DOI:** 10.1101/166470

**Authors:** José Guilherme Chaui-Berlinck, Luiz Henrique Alves Monteiro

## Abstract

The Frank-Starling Law of the heart is a filling-force mechanism, a positive relationship between the distension of a ventricular chamber and its force of ejection. The functioning of the cardiovascular system is usually described by means of two intersecting curves: the cardiac and vascular functions, the former related to the contractility of the heart and the latter related to the after-load imposed to the ventricle. The crossing of these functions is the so-called operation point, and the filling-force mechanism is supposed to play a stabilizing role for the short-term variations in the working of the system. In the present study, we analyze whether the filling-force mechanism is responsible for such a stability within two different settings: one-ventricle, as in fishes, and two-ventricle hearts, as in birds and mammals. Each setting was analyzed under two scenarios: presence of the filling-force mechanism and its absence. To approach the query, we linearized the region around an arbitrary operation point and put forward a dynamical system of differential equations to describe the relationship among volumes of ventricular chambers and volumes of vascular beds in face of blood flows governed by pressure differences between adjacent compartments. Our results show that the filling-force mechanism is not necessary to give stability to an operation point. The results indicate that the role of the filling-force mechanism is related to decrease the controlling effort over the circulatory system, to smooth out perturbations and to guarantee faster transitions among operation points.

**Summary Statement:** We address the role of the Frank-Starling mechanism and show that it has no role in the stability of the circulatory system. Rather, it accounts for decreasing the controlling effort and speeding up changes in cardiac output.

## Introduction

The nowadays-called Frank-Starling Law, or Heart Law, has a long history, being known since the beginning of 1830 (Katz, 2002). Such a “law” is a relationship between the filling of a ventricle and the force of contraction it develops (e.g., (Holubarsch et al., 1996)). In this way, it is also known as the heart filling-force relationship (Katz, 2002; Saks et al., 2006), the length-dependent activation (Solaro, 2007), or, even, the stretch-activation/calcium-activation (Campbell and Chandra, 2006). And despite the fact that fishes regulate cardiac output mainly by changes in stroke volume while mammals and birds control mainly heart rate, the filling-force mechanism (FFm) is found across all vertebrate classes (Shiels and White, 2008).

The relationship between length and force in the heart resembles the same relationship occurring in skeletal muscles. However, the steepness of the curve obtained for the heart suggested that beyond myofilament overlapping, there should be other, or others, mechanism involved in the phenomenon. Indeed, a calcium-activation process is fundamental for the increase in force due to an increase in length (e.g., (Moss and Fitzsimons, 2002; Niederer and Smith, 2009; Saks et al., 2004)). Be that as it may, it is important to note that the FFm is inherent to the heart cells themselves, without the participation of extrinsic controls as neural or hormonal ones. As stated in the opening of the review by Shiels and White (Shiels and White, 2008), “The Frank-Starling mechanism is an intrinsic property of all vertebrate cardiac tissue”.

Guyton and co-workers conceived an invaluable static approach to address the functioning of the cardiovascular system. We qualitatively illustrate this approach in Fig. 1A, where the abscises axis is the central venous pressure and the ordinate axis is the cardiac output. There, it can be seen two curves: the cardiac function (the ascending one) and the vascular function (the descending one).

**Figure 1.**
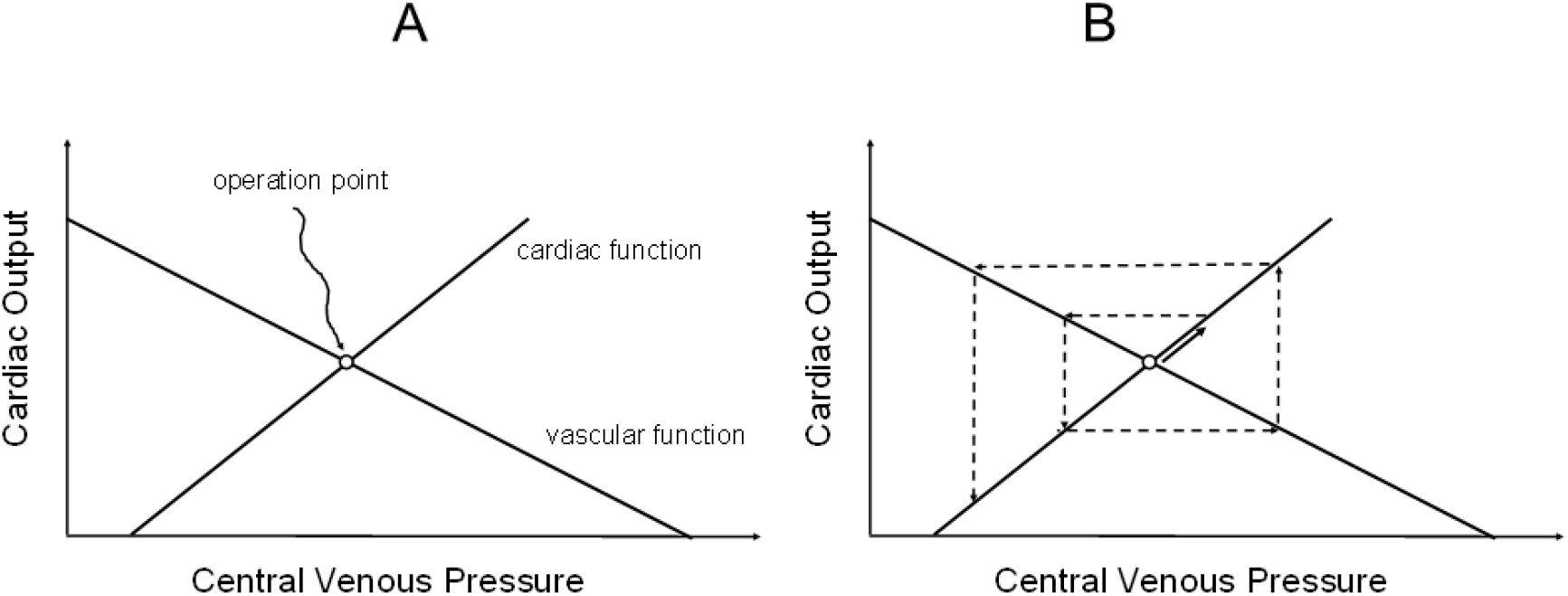
Cardiovascular operation point. **(A)** Usual representation of the cardiac and vascular functions resulting in an operation point of the heart. **(B)** Pictorial representation of a non-stable equilibrium (operation) point (an unstable focus in this case). The solid arrow represents an arbitrary perturbation from the operation point; the dashed lines represent a possible evolution path. This path is only for illustrative purposes and based on a cobwebbing approach of discrete dynamical systems.

The cardiac function ultimately represents the filling-force mechanism discussed above, since an increase in central venous pressure would elicit an increase in ventricular volume during the diastolic phase of cardiac cycle – which, in turn, would increase the contraction force resulting in an increase in cardiac output.

The vascular curve is, in fact, plotted the other way around as it is truly obtained (the experimental procedure is to cause changes in flow and measure the resulting pressure), and represents the dependence of central venous pressure in relation to blood flow (for details and insightful discussions of this subject, see (Brengelmann, 2003; Levy and Pappano, 2007)). The crossing of the two curves is the so-called operation point (OP) of the cardiovascular system.

Now, many textbooks and papers consider, implicit or explicitly, the OP as a stable equilibrium point, and that the FFm is responsible for such a stability. Let us give some examples.

–“… [OP] represent the stable values of cardiac output and central venous pressure at which the system tends to operate. Any perturbation … institutes a sequence of changes in cardiac output and venous pressure that restore these variables to their equilibrium values” ((Levy and Pappano, 2007), pg. 187).
–“[Frank-Starling mechanism] … applies in particular to the coordination of the output of the two ventricles. Because the ventricles beat at the same rate, the output of the two can be matched only by adjustments of the stroke volume.” ((Antoni, 1996), pg. 1814).
–“The heart maintains normal blood circulation under a wide range of workloads, a function governed by the Frank-Starling law” (Saks et al., 2006).
–“This important functional property of the heart supplies an essential regulatory mechanism by which cardiac output is intrinsically optimized relative to demand.”(Asnes et al., 2006).

Besides these citations, we can easily lengthen the list of those that, one way or another, consider the OP as an stable equilibrium point due to the FFm (e.g., (Fuchs and Smith, 2001; Moss and Fitzsimons, 2002; Niederer et al., 2011)).

As we see from the above-mentioned literature, students and physicians are lead to consider the filling-force mechanism as giving stability to the system.

However, if we take the (apparent) stability of the cardiovascular system as a *prima facie* evidence of the (supposed) stability generated by the FFm, we risk ourselves to fall in a circular reasoning. Actually, the OP could well be a neutral equilibrium point or, even worst, an unstable node or focus, all compatible with the curves that describe the OP (see Fig. 1B as an example). In effect, during undergraduate and graduate disciplines, one of us (JGCB) has trouble in explaining the stability of the OP from the vascular and cardiac curves. If one examines with care the diagram, a perturbation in the OP would not be dampened in the following cycle(s) but instead, it would be amplified.

Why does this occur? Because the OP-diagram is not a diagram concerning the dynamical phase-space of the variables. It shows a static 2D relationship between a pair of variables that belong to a higher dimensional space: the curves are somehow projections of the null-clines of the whole system (note: in the case of one-ventricle hearts, as it will be also modelled, the OP-diagram is a construct from a lower dimensional space, but this is not really important here).

In plain English, the OP-diagram does not, and cannot, reveal how changes in one variable (say left cardiac output) alters the other (say central pulmonary venous pressure) because there are missing variables. If the vascular curve refers to the vena cava, then the cardiac curve should be for the right ventricle. If the vascular curve refers to the pulmonary veins, then the cardiac curve should be for the left ventricle. However, as usually presented, the OP diagram mixes up the two sides of the heart. Once we recognize this, we understand that, for two-ventricle hearts, one needs four state variables to compose the whole picture (despite this obviously prevents a 2D representation): the systemic pressure, the right ventricle output, the pulmonary pressure and the left ventricle output. Therefore, there are two operation points: one for the left side and one for the right side of the heart.

In a more formal language, the diagram of the vascular and of the cardiac curves (Fig. 1) as obtained does not have an associated vector field in the phase-space that represents the possible trajectories of the system given a perturbation from the OP. Thus, the conundrum is whether the OP is a stable equilibrium point due to the filling-force mechanism, which, in the end, guaranties that both beat-to-beat variation and the matching between the ventricles can be sustained *without any regulatory loop extrinsic to the heart*.

The filling-force mechanism is found among all vertebrate classes, as stated in before. However, many vertebrates have single-ventricle hearts, and so, there is no match necessities between the outputs of two ventricles beating simultaneously. Moreover, exactly these vertebrates belong to the predecessor lines of the two-ventricle hearts of mammals, birds and some reptiles. Thus, in evolutive terms, the FFm precedes output-matching necessities.

Fishes regulate cardiac output mainly by systolic volume and it is considered that the FFm is responsible for the adjustment of ejection in face of large changes in ventricle volume (Shiels and White, 2008). The ascending limb of the relationship between developed tension and sarcomere length is much broader in these animals than in mammals and birds, indicating a wider range of adequate ventricular pressure responses in face of increases in chamber volume (Shiels and White, 2008). Despite the fact that these considerations seem to address the question of the stability of a given equilibrium point in fishes, in fact they are related to the transitions among operating points governed by a series of systemic changes (e.g., changes in metabolic demand, muscle contraction, autonomic tonus, etc.). Counterintuitively as it may sound, the latter, transitions, does not imply the former, stability, indeed.

The present study aims to answer the questions of the role of the filling-force mechanism in the stability of an operation point and of the role of the FFm in output-matching. These questions are approached by the analysis of a dynamical system representing the acute and intrinsic coupling between cardiac output and central venous pressure. We analyze two settings of this coupling, one concerning the single ventricle system of fishes and the other concerning the two-ventricle system of mammals, birds and some reptiles. The settings are analyzed in two different scenarios: (A) the filling-force mechanism actuating in the ventricular chamber; and (B) a fixed force is exerted by a ventricular chamber. These two scenarios are intended to allow for a comparison of what would happen if the FFm were absent and so, to answer the proposed questions.

## Preliminary considerations

### Mechanistic description and cardiac dynamics

The functioning of the cardiovascular system is governed by a set of variables. This set includes vascular capacitances, vascular impedances, blood rheology, total blood volume, autonomic nervous system tonus (e.g., (Holubarsch et al., 1996; Hoppensteadt and Peskin, 2002)). For the purposes of the present analysis, these variables would be considered as constants during the timeframe of interest. This defines what is meant by “acute” and by “intrinsic” that we put above. In other words, we are saying that there is more than one time scale to describe the system, and we shall investigate one that operates at a rate compatible of a heartbeat interval. In doing so, we are lead to consider that in the vicinities of an OP the system behaves linearly.

In this instance, the total volume of fluid (explicitly, blood), VT, is constant and equals the sum of the volumes in each compartment j of the system:

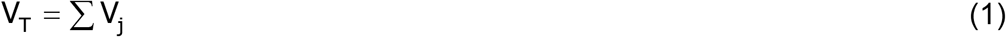

We use the Hagen-Poiseuille model to describe flow between two points i and j of the circulatory system:

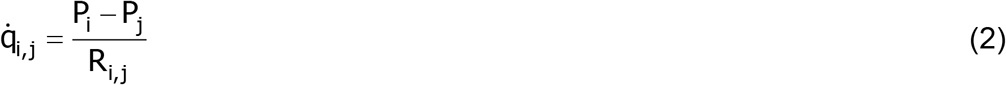

In which q is the flow between compartments i and j, P is the pressure in a given compartment and R is the resistance imposed to the flow between the compartments. Notice that the resistance term encloses physical constants of the system, such as mean radius and length of the vessels, viscosity of the fluid, etc.

The pressure in a given compartment j is the volume V of blood present in the compartment divided by the capacitance β of the compartment (here we consider the capacitance as a constant in the small range of volume variations we analyze):

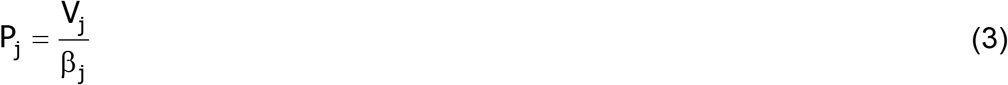

Eqn 1-3 form the core of the subsequent models in which the time variation in the volume of a given compartment j is the result of the inflow and outflow of blood:

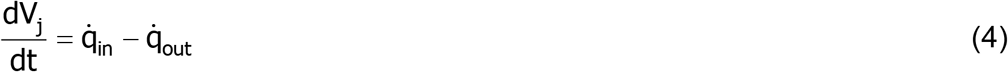

Since total volume is constant, then follows that:

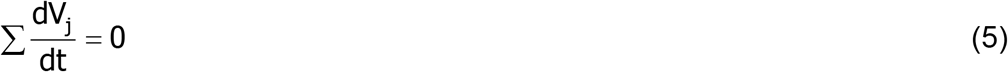

As stated before, the timeframe of reference is related to a heartbeat, which is composed by two phases. During systole, the heart ejects but does not receive blood. During diastole, the reverse is true. Therefore, when we employ Eqn 4 we are referring to mean values during the cardiac cycle. To incorporate such a cycle in the mean-valued model, we consider that, during diastole, the capacitance of the ventricle tends to infinity, and, therefore, the circulatory tree fills the heart against a near-zero pressure. During the systole, the ventricle develops a certain pressure (force), and this pressure is related to the volume of the ventricle. This is the filling-force mechanism, indeed.

The model is intended to study the behavior of the system near an operation point. Therefore, we employ a simple positive linear relationship between volume and pressure (force). This means that we are neither modeling any transition between two distant operation points nor pathological conditions where the FFm might be inverted (i.e., the greater the ventricular volume the lower the developed force).

## Modeling and Results

### From fish … One-ventricle hearts

Let the indexes H represent the heart chamber and S the vascular tree, respectively (Fig. 2).

**Figure 2.**
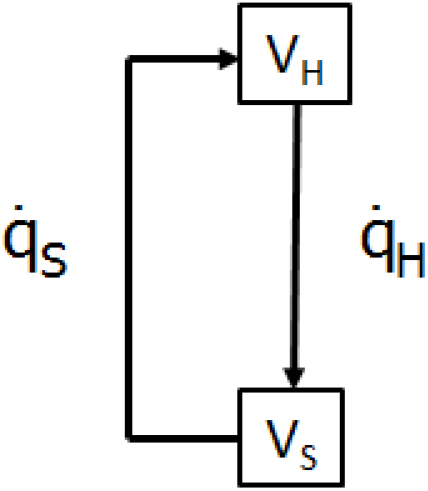
Schematics of the model of the one-ventricle heart system. The state variables heart volume (VH) and systemic volume (VS) are in boxes. The arrows indicate blood flows.

#### Scenario (A): the filling-force mechanism actuating in the ventricular chamber

The outflow from the heart (inflow to the vascular tree) and the outflow from the vascular tree (inflow to the heart) are:

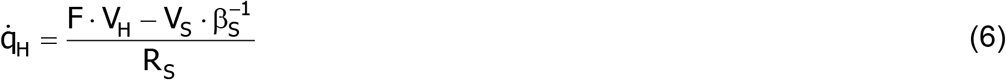

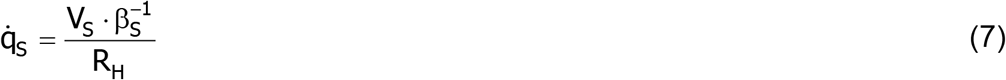

In which F is the linear coefficient of the relationship between ventricle volume and developed pressure (the filling-force mechanism). For the sake of notation, we define:

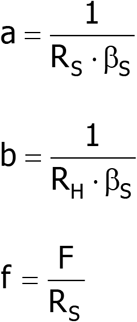

Coefficients a, b and f have units of [pressure] · [volume]^-1^ · [resistance]^-1^. Since resistance to flow have units of [time] · [pressure] · [volume]^-1^, the coefficients end up as [time]^-1^ (i.e., inverse of time-constants).

Because the time variation in total blood volume is zero (Eqn 5), then, from Eqn 4, the system is described by the following differential equation:

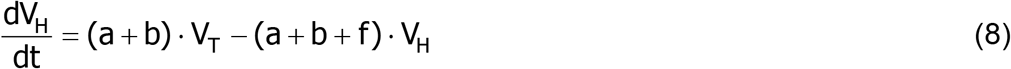

By equating dVH/dt to zero, we obtain the value of the cardiac volume (and, consequently, the one of the vascular tree as well) at the equilibrium point of the system, denoted by an “*”:

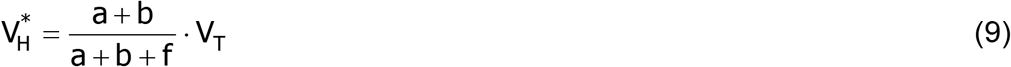

In fact, Eqn 8 can be integrated straightway and we have:

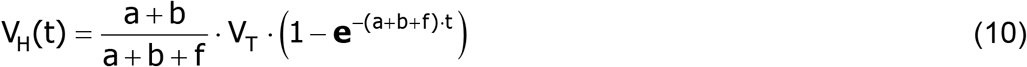

In which **e** is the base of the natural logarithm.

#### Scenario (B): a fixed-force is exerted by a ventricular chamber

We use the subscript “k” to indicate the parameters and the variables in this fixed-force scenario. The outflow from the heart (inflow to the vascular tree) becomes:

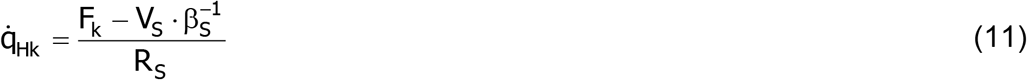

In which Fk is the fixed-force term. The outflow from the vascular tree (inflow to the heart) remains the same as in Eqn 6. The differential equation describing the dynamics of the system is now:

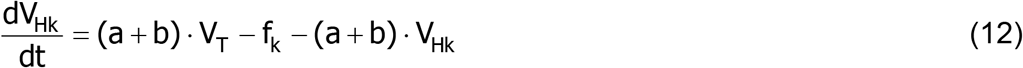

Notice that the constant fk has units of [volume] · [time]^-1^, i.e., flow. By integrating Eqn 12 results in:

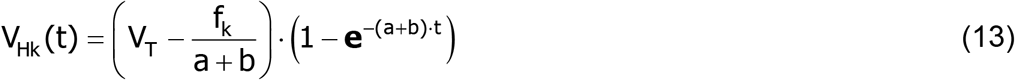

And the value of the cardiac volume at the equilibrium point is:

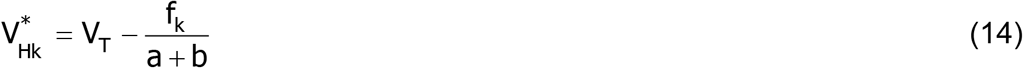

Eqn 14 shows that, if the fixed-force term (represented by fk) is much greater than the sum of a + b, the heart chamber would become completely empty of blood.

#### Stability of the Equilibrium Point

Both Eqn 10 and 13 reveal that their respective equilibrium points are an asymptotically stable node: both eigenvalues are negative real numbers (e.g.,(Monteiro, 2011)). Therefore, irrespectively to the presence of the FFm, the one-ventricle circulatory system has a stable operation point.

### … to philosopher ^1^ - Two-ventricle hearts

As stated in the Introduction, we need four state-variables to describe the two- ventricle hearts: left ventricle (L), systemic vascular bed (S), right ventricle (R) and pulmonary vascular bed (G – we use G for “Gas exchanger organ” instead of “P” that would cause confusion with pressure). See Fig. 3.

**Figure 3.**
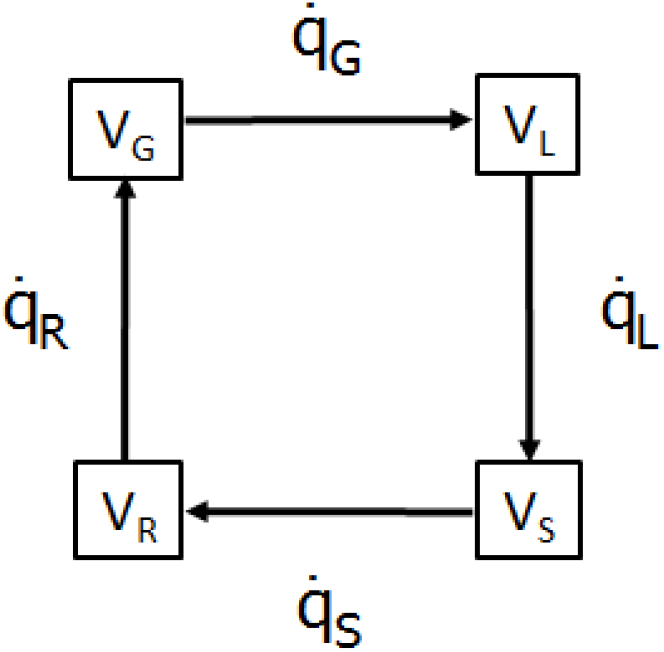
Schematics of the model of the two-ventricles heart system. The state variables left ventricle volume (VL), systemic circulation volume (VS), right ventricle volume (VR) and gas-exchanger circulation volume (VG) are in boxes. The arrows indicate blood flows.

#### Scenario (A): the filling-force mechanism actuating in the ventricular chamber

Flows are given by the following equations:

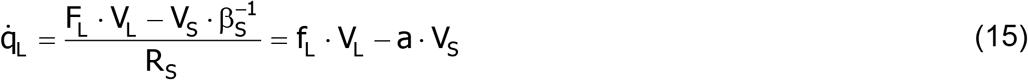

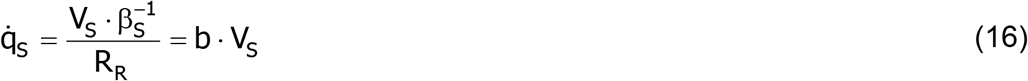

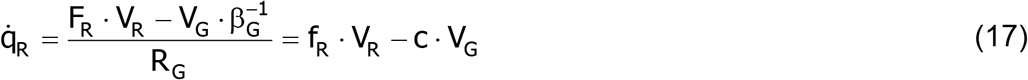

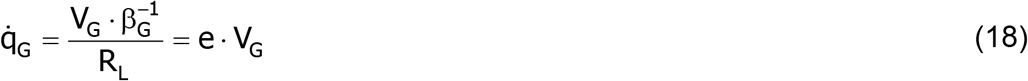

In which we employ the same short notation as in the preceding section for the sake of clarity. From the equations of flow and Eqn 5, we have the following set of coupled differential equations to describe the system:

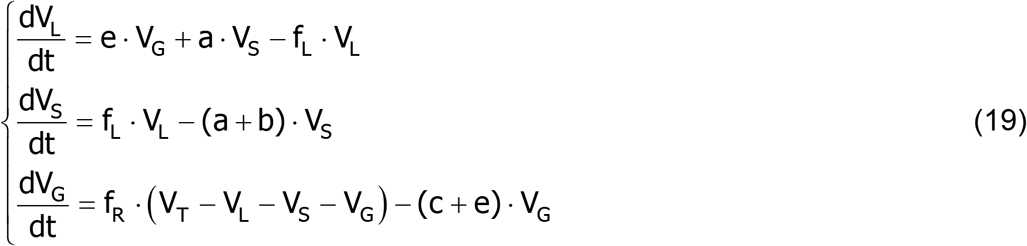

The volumes at the equilibrium point of the system are (we let VS* and VG* as functions of VL*):

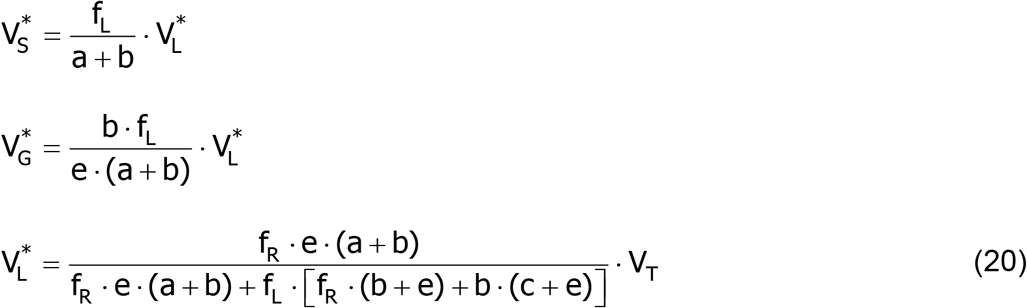

Just to check the feasibility of Eqn 20, if fR = 0, i.e., the right ventricle has no ejecting force at all, then the whole volume of blood would be retained in the right ventricle, while if fL = 0, then the volume is completely retained in the left ventricle. If both fR and fL go to zero simultaneously, then one has a proportion of blood retained in the right side and other in the left side, as in stagnation conditions. These extreme results are in accordance with what one would anticipate within this simplified framework of the circulatory system.

#### Stability of the equilibrium point in the presence of the filling-force mechanism

The stability of the equilibrium point is given by setting the determinant of the Jacobian of the system to zero:

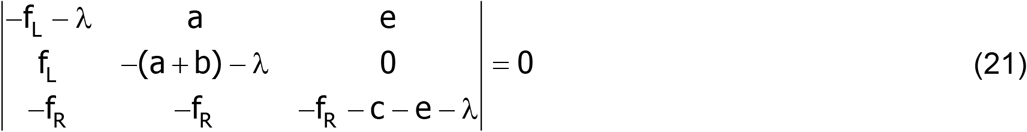

In which λ is an eigenvalue of the system. This determinant corresponds to the following characteristic equation:

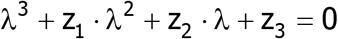

The coefficients zi are:

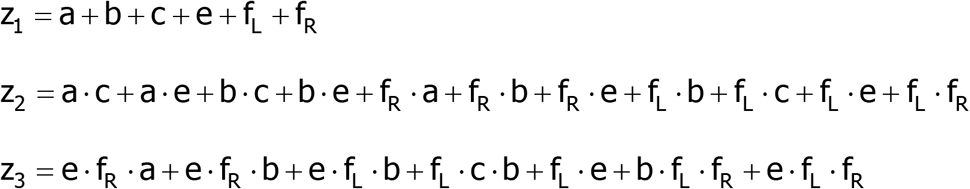

For the equilibrium point be asymptotically stable, the following conditions must be satisfied:

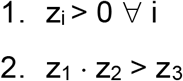

Since all parameters are positive, condition 1 is satisfied. Plain inspection of the coefficients shows that condition 2 is also satisfied. Therefore, the equilibrium point of a two-ventricle system in the presence of the filling-force mechanism is asymptotically stable.

#### Scenario (B): a fixed-force is exerted by a ventricular chamber

The system is described by the following coupled differential equations, where the subscript k indicates the fixed force:

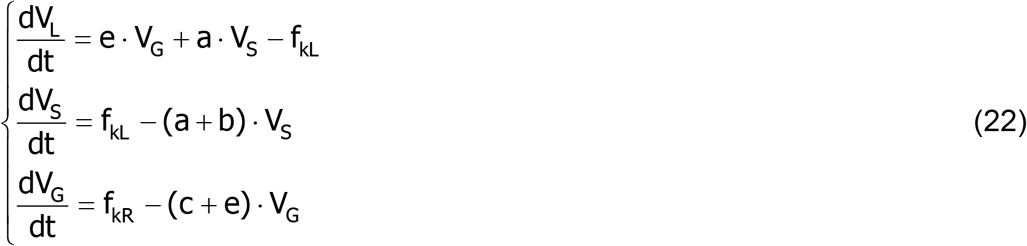

The volumes of the compartments S and G at the equilibrium point of the system are:

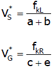

From these values in the equation of dVL/dt, we obtain that the following relationship must hold in order to the system have an equilibrium point:

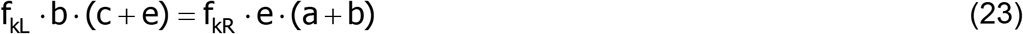

Therefore, unless condition 23 is fulfilled, the system will not attain an equilibrium point at all. Also notice that the volumes of two compartments are not obtained (see below – in this case, these volumes are from the left and the right ventricles, but this due to the form that we delineate system 22 – the relevant point is that there are two unknown volumes).

#### Stability of the equilibrium point in the presence of a fixed-force of ejection

We obtain the following determinant of the Jacobian of the system 22:

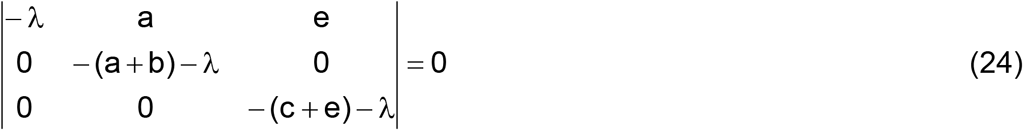

Therefore, the system has an asymptotically stable subspace with two real eigenvalues (λ_1_ = -(a + b) and λ_2_ = -(c + e)) and a central manifold corresponding to λ_3_ = 0. This central manifold represents the indeterminacy of the two volumes (VL and VR in this case). Let VH = VL + VR. Since:

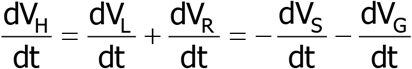

The system becomes simply:

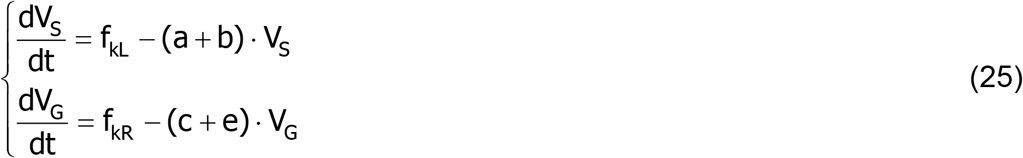

In a very similar way of what happens in the case of the one-ventricle hearts, the system is asymptotically stable even in the absence of the filling-force mechanism and, considering condition (23), one way to write the heart volume is:

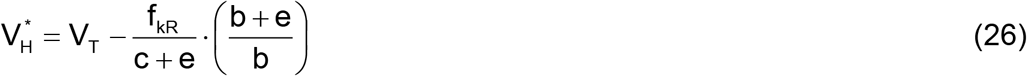

## Discussion

The stability of the operation point of the cardiovascular system is usually taken for granted as a result of the Frank-Starling Law, i.e., the filling-force mechanism of the heart. However, the OP diagram does not convey sufficient information to conclude that such an intrinsic mechanism of the myocardium truly would bring up stability to the system in a beat-to-beat basis.

In the present study, we approach this question by investigating the behavior of a dynamical system, representing a circulatory system, in the vicinity of an operation point. In such a vicinity, the temporal variation of a set of relevant physical variables in the cardiovascular system is taken as null, i.e., we investigate the behavior of the system within a fast time scale, roughly corresponding to the heartbeat interval. In this sense, all the sympathovagal inputs to the heart are considered as constants, as well as changes in blood volume, rheological factors, etc.

The first important conclusion of the study is that both types of circulatory systems, i.e., one-ventricle and two-ventricle hearts, are asymptotically stable even in the absence of the filling-force mechanism. In other words, if a given operation point exists, it is stable, and the system will return to such an OP after suffering a perturbation, irrespectively of the presence of the FFm (and without any extrinsic regulatory loop).

Therefore, the question now becomes more inclusive, since one has to understand the role of the filling-force mechanism without evoking its alleged and putative responsibility in stabilizing the operation point.

Due to the similar results between the systems with one and two ventricles, let us focus in the one-ventricle heart for simplicity. Eqn 9 and 14 describe the volume in the heart compartment for one system with and for another one without the filling-force mechanism, respectively. Fig. 4 shows a plot of these functions (the 5% volume line is indicated simply as a reference to a usual value of the volume in the heart in relation to the volume of blood).

**Figure 4.**
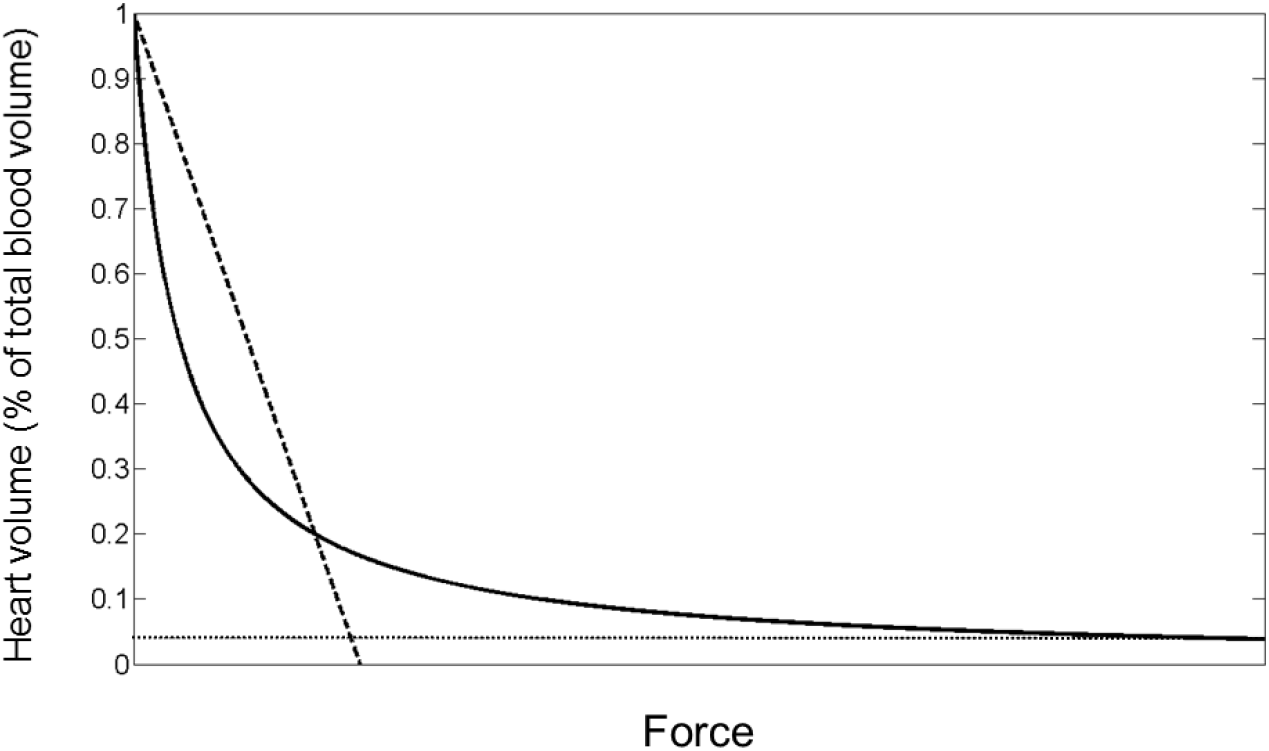
Comparison between the effects of varying the force terms in the two different scenarios analyzed. (Eqn 9 and 14). The y-axis represents the fraction of blood in the cardiac chamber in relation to total blood volume. The x-axis represents force, i.e., the terms f and fk (it must be kept in mind that f and fK have different dimensions). Continuous line: volume in the scenario with the filling-force mechanism. Dashed line: volume in the scenario with a fixed-force exerted by the ventricular chamber. Dotted line: 5% of total blood volume. The sum of the terms a and b in both Eqn 9 and 14 is 1 for the simulations shown in the plot.

Despite the risk of becoming repetitive, let us put it once again: both scenarios allow for the existence of stable OPs. In addition, as already stated (see Results), if the force term tends to zero, the total blood volume tends to be retained in the cardiac chamber (left-hand side in Fig. 4). In the vicinities of the zero-force, the heart volume of the system with the filling-force mechanism shows a steeper relationship with force than the fixed-force system. However, from a certain volume down, the linear relationship of the fixed-force becomes steeper than the asymptote of the filling-force mechanism system. Thus, close to the range of reasonable heart volumes, the fixed-force system shows a higher variation in the volumes of its compartments in face of variations in force, while the filling-force system has a smooth response.

Controlling effort (e.g. (Kirk, 2012 pg. 259; Todorov and Jordan, 2002)) and computational complexity (e.g. (Benenti, G. Casati,G. Strini, 2007 pg. 24; Moller and Smolka, 1965)) are somehow related to energy waste and resources allocation by the controller system or the resolution algorithm in a given task. Considering that the resistances, capacitances and even the myocardial force itself (irrespectively to the scenario) are under adjustments regulated by the autonomous nervous system, the smoothness brought by the filling-force mechanism ends up as a lower effort on the controller unit (i.e., lower energy demand and/or use of system resources).

Inspection of Eqn 20 shows that the controller unit can operate a variation in one given parameter (say, systemic resistance in the coefficient “a”) and the circulatory system will self-adjust its volumes accordingly. On the other hand, in the scenario with fixed-force terms, inspection of Eqn 23 shows that the controller unit must operate simultaneous variations in at least two parameters in order to guarantee the working of the system.

Thus, the second conclusion we can draw is that the filling-force mechanism has a role in decreasing the controlling effort external to the circulatory system (note that this has nothing to do with the stability of an operation point discussed above). The absence of the FFm does not preclude variations to be operated in the circulation, but the presence of the FFm smooths out perturbations more easily.

Then, the next inevitable question is whether the filling-force mechanism plays some role in heart rate variability. Heart rate suffers variations on a beat-to-beat basis. The most prominent are changes associate to ventilation (respiratory sinus arrhythmia), but many other factors are also interconnected to these variations, resulting in a multifaceted composition of frequencies. The beat-to-beat modulation of heart rate is due to a number of feedback loops that end up through a common dual efferent path, the sympathetic and parasympathetic branches of the autonomic nervous system (e.g., (Aubert et al., 2003; Stauss, 2003)). Also, there might exist some intrinsic innervation in the heart itself whose role is not well established (Stauss, 2003). This modulation gives rise to the so-called “heart rate variability”, and such a variability is an important sign of an adequate functioning of the cardiovascular system (e.g., (Stauss, 2003; TASK FORCE, 1996)).

In this sense, the third relevant conclusion of the present study comes from the inspection of the eigenvalues of a system with the filling-force mechanism and of a similar system (i.e., a system with the same set of values for the parameters of the vascular bed) with a fixed ejection force. For the one-ventricle hearts, this can be directly evaluated in Eqn 10 and 13 for the cases with the FFm and without it, respectively. Considering that the volume of blood in the heart is approximately 5% of the total blood volume, from Eqn 9 we obtain that the filling-force term would be roughly 19-fold greater than the sum of the other two terms, a and b. This results in a returning to the operation point twenty times faster in the presence of the filling-force mechanism than in its absence.

For the two-ventricle hearts without the FFm, the eigenvalues of the stable sub-space are shown in Eqn 24. Although we did not directly compute the eigenvalues of two-ventricle hearts when the filling-force mechanism is present, we can have a glimpse of what occurs in them. Because the sum of the eigenvalues of a system equals the trace of the Jacobian matrix, then we can observe that both terms fL and fR, related to the filling-force mechanism, take part in at least one of the eigenvalues of the system (see Eqn 21). Therefore, similarly to what happens in the one-ventricle hearts, two-ventricle systems will also return to the operation point faster in the presence of the filling-force mechanism than in its absence.

Thus, our third conclusion is in regard of the time-constant of a system: the filling-force mechanism allows for a much faster return to an operation point after a perturbation. In other words, despite the fact that an existing operation point is stable even in the absence of the FFm, its presence guaranties the operation point to be regained in a fraction of the time than if there were no such a mechanism.

Heuristically, we might consider that when the system transits from a previous operation point to a new one, the former is a perturbation in relation to the latter (notice that this is not the mathematical definition of “perturbation”). In this sense, the transition among operation points would be speeded up by the filling-force mechanism. In a similar line of reasoning, this speeding up potentially contributes to non-autonomic components of heart rate variability, particularly in the high-frequency range.

In conclusion, differently from what is currently held, the filling-force mechanism is not necessary in order to give stability to an operation point in a circulatory system, whether composed by a heart with a single or with two ventricles. Our modelling supports that the role of the filling-force mechanism is related to decrease the controlling effort over the circulatory system, to smooth out perturbations and to guarantee faster transitions among operation points.

## Footnotes

1 “From Fish to Philosopher” is a classical book by Homer William Smith (1959).

## Acknowledgements

JGCB is grateful to his graduate students who discussed a preliminary version of the mathematical model.

## Supporting Data and Data Availability

This study has no supporting material or data.

## List of Symbols and Abbreviations

**Symbol or Abbreviation**
FFm: filling-force mechanism
OP: operation point
V: blood volume
P: pressure
R: resistance
β: capacitance
q: flow
F: coefficient of force

**subscripts**
T: total
j: a general compartment
k: fixed-force scenario
H: one-ventricle chamber
S: systemic vascular bed
G: pulmonary vascular bed
L: left ventricle
R: right ventricle

